# Improving protein secondary structure prediction by deep language models and transformer networks

**DOI:** 10.1101/2022.11.21.517442

**Authors:** Tianqi Wu, Weihang Cheng, Jianlin Cheng

## Abstract

Protein secondary structure prediction is useful for many applications. It can be considered a language translation problem, i.e., translating a sequence of 20 different amino acids into a sequence of secondary structure symbols (e.g., alpha helix, beta strand, and coil). Here, we develop a novel protein secondary structure predictor called TransPross based on the transformer network and attention mechanism widely used in natural language processing to directly extract the evolutionary information from the protein language (i.e., raw multiple sequence alignment (MSA) of a protein) to predict the secondary structure. The method is different from traditional methods that first generate a MSA and then calculate expert-curated statistical profiles from the MSA as input. The attention mechnism used by TransPross can effectively capture long-range residue-residue interactions in protein sequences to predict secondary structures. Benchmarked on several datasets, TransPross outperforms the state-of-art methods. Moreover, our experiment shows that the prediction accuracy of TransPross positively correlates with the depth of MSAs and it is able to achieve the average prediction accuracy (i.e., Q3 score) above 80% for hard targets with few homologous sequences in their MSAs. TransPross is freely available at https://github.com/BioinfoMachineLearning/TransPro

## 1 Introduction

Studying protein structures and their functions is critical for addressing many biomedical problems such as protein engineering and drug design[1–4]. Predicting the protein secondary structure from sequence can provide useful information for studying tertiary and quaternary structures of proteins and their function [5–7].

Protein secondary structure (SS) can be defined as the local configuration of the polypeptide chain. The two regular local configuration types, alpha-helix (H) and beta-strand (E), determined by the hydrogen bonding patterns, were initially proposed in 1951[8, 9]. The non-regular local configuration is often called coil or loop (C). The annotation of the protein secondary structure is further expanded to the eight types and is commonly computed from the tertiary structure of proteins by the standard annotation tools such as DSSP[10]. For simplification, the 8-state representation is usually reduced to the 3-state representation (alpha-helix (H), beta-strand (E), and coil (C)). Since the tertiary structures of most proteins are unknown, there is a need to directly predict the protein secondary structure from sequence.

Many computational protein SS prediction methods have been developed over the last several decades. One major improvement in the protein SS prediction came from the innovation in enhancing the input information. Instead of using a single sequence of a target protein to predict its secondary structure, the sequence profile-based methods[11–19] compute the sequence profile from protein multiple sequence alignment(MSA) of the protein[20–23] to extract the evolutionary information for SS prediction. Using protein sequence profile for secondary structure prediction is substantially more accurate than using the single sequence of a target protein. The sequence profile is usually represented by expert-designed statistical models such as position-specific scoring matrices[21] and hidden Markov models[22]. As more and more protein sequences are produced, reasonable multiple sequence alignments can be generated for most proteins to construct good sequence profiles, the profile-based secondary structure prediction methods have reached a high accuracy (e.g., *>*80%) on most benchmarks. However, for some proteins that do not have many homologous sequences in the protein sequence databases for constructing reasonable multiple sequence alignments and profiles, the profile-based prediction methods cannot deliver optimal results. In this case, the single-sequence based prediction method that only depends on the single sequence information itself[18, 24] has been developed to improve the prediction accuracy.

Since 1990s, neural networks had been shown to be the most successful methods for protein secondary structure prediction [11, 13]. When deep neural networks consisting of many more layers than traditional shallow neural networks started to achieve significant success in several computing domains such as image processing and computer vision in mid-2020s, different deep learning architectures such as deep belief networks, gated recurrent neural networks, long- and short-term memory networks, convolutional neural networks, residual networks, and inception networks have been applied to the secondary structure prediction problem [12, 25–27].

Recently, the language models in the natural language processing (NLP) were adapted for protein sequence analysis[28–31], creating a new avenue to improve protein secondary structure prediction. The language models that can interpret the meaning of the words in a long-range context is a promising technique to handle the long-range amino acid interactions in protein sequences that are needed for predicting some secondary structures in proteins, particularly beta-sheets that often involve the long-range interactions. In this work, we explore the application of language models in protein SS prediction and develop a predictor - TransPross - based on the transformer network and the attention mechanism to effectively obtain evolutionary information from the raw MSA directly to predict secondary structure. Tested on several different test sets including the TransPross test set, CASP13 free modeling (FM) test set and CASP14 test set, TransPross achieves better performance than the state of the art of the sequence profile-based methods. The analysis of the performance of TransPross on hard targets with few homologous sequences in their MSAs also shows it often works well even when the MSAs are shallow. By taking a MSA as input, TransPross can be readily applied to many proteins that users may have obtained MSAs for. It can also be combined with any MSA generation tool to predict secondary structure prediction, which is more flexible than most traditional secondary structure predictions that use their own built-in MSA generation and profile generation programs to prepare the input. Therefore, TransPross is a useful tool that complements the existing protein secondary structure prediction tools.

## 2 Results

### 2.1 Comparison with single-sequence-based methods and profile-based methods

We compare TransPross with one of the most widely used state-of-the-art protein secondary structure prediction method - PSIPRED on three different test sets (TransPross test set,CASP13-FM test set consisting of the free modeling (FM) targets of the 13th Critical Assessment of Techniques for Protein Structure Prediction (CASP13) and CASP14 regular full-length targets with sequence length ≤500. FM targets were considered hard targets because they did not have similar structures in the Protein Data Bank (PDB) when they were released. PSIPRED can predict secondary structures from either sequence profile or a single protein sequence. PSIPRED-profile accepts the sequence profile calculated from multiple sequence alignments as input. PSIPRED-single predicts secondary structure from a protein single sequence. In addition to comparing TransPross with PSIPRED-profile, we also use PSIPRED-single and another single-sequence-based method SPOT-1D as the baseline to evaluate TransPross. The performance of the methods is evaluated by the Q3 accuracy score, which is the percent of residues of the proteins in a dataset that are correctly assigned into three categories (i.e., helix, beta-strand, and coil). As the results shown in Table 1 (a) and (b), TransPross substantially outperforms the single sequence-based methods(SPOT-1D, PSIPRED-single) as expected. The Q3 accuracy score of TransPross for the three test sets are 84.42%, 81.04% and 80.71% respectively, which is also better than that of a state-of-the-art profile-based method PSIPRED-profile.

**Table 1:** Comparison of TransPross with other predictors in terms of the Q3 accuracy score on different test sets: (a)TransPross test set and CASP13 FM test set (b) CASP14 test set

| Method | TransPross-test | CASP13 |
| --- | --- | --- |
| TransPross | **84.42** | **81.04** |
| SPOT-1D | 76.97 | 73.21 |
| PSIPRED-single | 71.03 | 68.05 |
| PSIPRED-profile | 82.88 | 80.60 |

| Method | CASP14 |
| --- | --- |
| TransPross | **80.71** |
| PSIPRED-profile | 79.62 |

(b) Comparison of TransPross with PSIPRED-profile on CASP14 test set.

### 2.2 Effect of the quality of MSAs on prediction accuracy

To evaluate the impact of the quality of multiple sequence alignments (MSAs) on the performance of TransPross, we test TransPross on two different types of inputs: MSAs generated by using HHblits to search protein sequence against the big fantastic protein sequence database (BFD)[32], and MSAs generated by using DeepMSA[33] to search against Uniref30 and the mgnify databases. The quality of a MSA is determined by multiple factors such as the alignment accuracy, the number of homologous or non-homologous sequences in the MSA, and the redundancy in the MSA. Here, we simply calculate the number of effective sequences(Neff) in the MSAs to approximate their quality [34]. In general, the higher an Neff, more diverse and informative a MSA is. The average Neff for the two different inputs for 17 CASP13-FM targets is shown in Table 2, MSA BFD denotes the MSAs generated from the BFD database, while MSA DeepMSA represents the MSAs generated by DeepMSA. Because DeepMSA searches against several databases, MSA DeepMSA has a higher Neff than MSA BFD in most cases.

**Table 2:** The effect of Neff on the prediction accuracy of three-state secondary structure on 17 CASP13 FM domain targets.

| Input | Q3(%) | Neff |
| --- | --- | --- |
| MSA_BFD | 76.09 | 202 |
| MSA_DeepMSA | 81.04 | 382 |

In Table 2, there is a positive relationship between the average Q3 accuracy score of TransPross and the average Neff of MSA on the 17 CASP13 FM targets. Figure 1 plots the Q3 score against Neff of the 17 CASP13 FM targets. The results indicate that increasing the number of effective sequences in MSAs is important for improving secondary structure prediction, which is similar to protein contact prediction [34] and tertiary structure prediction [35]. The Pearson’s correlation coefficient between the per-target Neff and the corresponding Q3 accuracy score on the 17 CASP13 FM targets is 0.52.

**Fig. 1:**
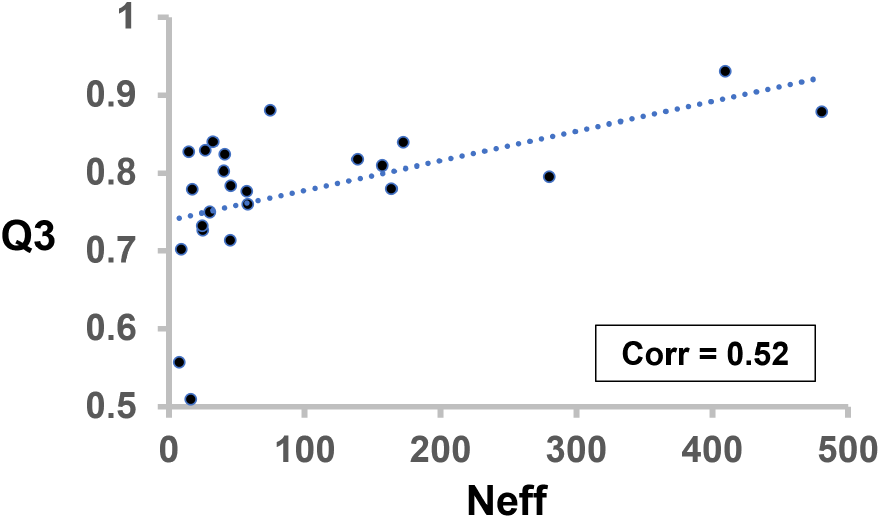
The scatter plot of the Q3 accuracy score of secondary structure prediction versus Neff with linear fitting curve (Pearson’s correlation coefficient = 0.52)

### 2.3 Secondary structure prediction for hard targets with shallow MSAs

Targets with relatively few sequence homologs in their MSAs are generally defined as hard targets and the accurate predictions for them are likely more helpful for studying their structures and function. We assess the performance of TransPross on the 12 CASP13 FM targets with shallow MSAs (Neff *<* 50). Interestingly, TransPross is able to achieve a pretty good average Q3 accuracy sore of 81.25%, slightly higher than PSIPRED-profile’s 81.22%. Figure 2 illustrates a head-to-head comparison of TransPross predictions with the PSIPRED-profile predictions on 12 CASP13 FM targets with shallow alignments(Neff *<* 50). In total, TransPross performed better in 8 of 12 cases, demonstrating its capability of accurately predicting secondary structure for hard targets with a small number of sequence homologs.

**Fig. 2:**
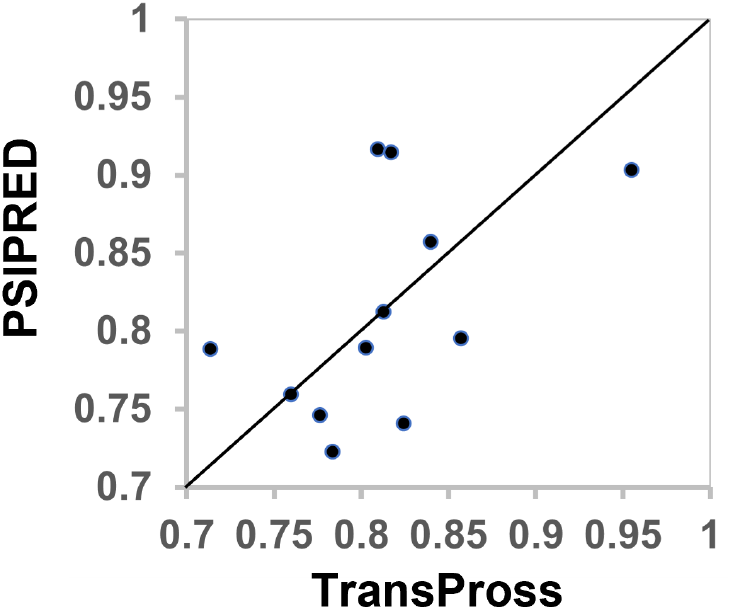
Performance of TransPross versus PSIPRED-profile on 12 CASP13 FM domain targets with shallow MSAs (Neff *<* 50) in terms of the Q3 accuracy score (denoted by a dot in the plot)

### 2.4 Running time

We empirically calculated the running time of TransPross on the CASP13 FM targets with an average length of 148 residues. If the MSA of a target is provided, TransPross typically requires about 30 seconds on average to complete the prediction with a single Tesla V100 GPU. Because MSAs are often available for many proteins in advance, TransPross can be applied to them to quickly generate secondary structure predictions. If MSAs are not available, users can use any MSA generation tool or the programs in the TransPross package to generate MSA first, which is generally much slower than predicting secondary structure from MSA with TransPross.

## 3 Materials and Methods

### 3.1 Datasets

We selected protein targets deposited into the Protein Data Bank(PDB) before May 2019 and extracted their true secondary structures. After filtering out the redundant sequences at the sequence identity cutoff of 90% by MMseqs2[36] and setting the sequence length within the range [50, 500], there are 36,334 proteins left. We first randomly selected 5% of the targets to create the TransPross test set and the rest of the targets were used for training and validation. Apart from the TransPross test set, the other two test sets are 17 CASP13 FM domains and 40 CASP14 regular targets with length ≤500. To ensure there was no overlap between the training data and test sets, we further filtered out the sequence redundancy between the training data and all the three test sets at the sequence identify cutoff 30% by using MMseqs2[36]. We extracted the true 3-state secondary structures of the proteins (i.e., strand E, helix H, and loop C) from their true tertiary structures as labels using DSSP[10]. The objective is to predict the secondary structure label for each residue of a protein from its sequence.

### 3.2 Protein sequence language model and transformer architecture

We apply a transformer network with the attention mechanism[37] that has achieved success in natural language translation to detect the relevant sequence context across the entire protein sequence for each amino acid position to predict its secondary structure type. The deep learning architecture is useful to capture long-range interaction between amino acids (like words in an English sentence) relevant to secondary structure prediction. The customized protein sequence-to-secondary-structure translation model of TransPross shown in Figure 3 adopts an encoder-decoder structure. The encoder maps an input MSA of a protein in the symbolic representation to a sequence of continuous values (internal features) with the dimension of *L ×* 512(L: the protein sequence length). From the internal features, the decoder then generates an output sequence of 3-state protein secondary structures one by one in an auto-regressive way[38].

**Fig. 3:**
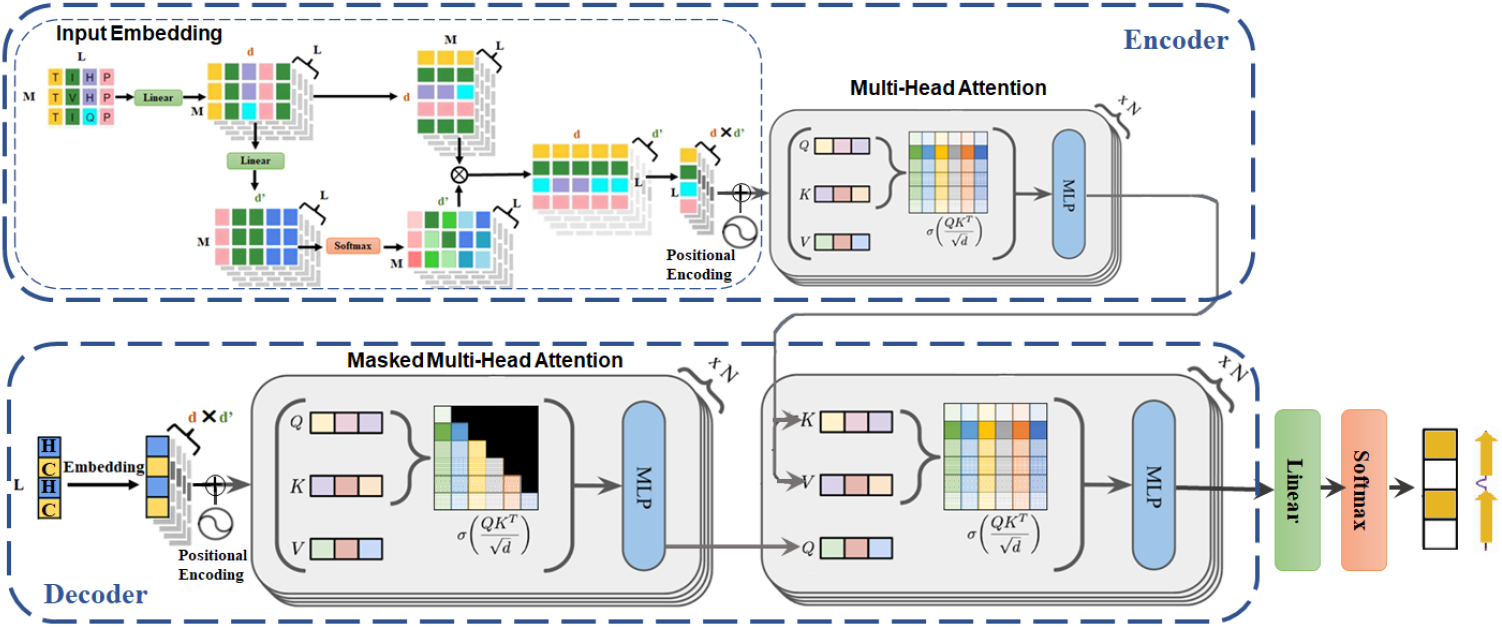
The architecture of TransPross. It has two main modules: an encoder and a decoder. The input for the encoder is the MSA of a protein (M). The input for the decoder includes both the secondary structures prior to the current position under consideration and the output of the encoder. From the input, the decoder predicts the three-class secondary structure for each position in the protein sequence.

Given the MSA for a target protein as the input, TransPross uses the encoder-decoder process to generate the secondary structure of the protein.The encoder consists of the input embedding component and self-attention component. The input embedding component first converts the input tokens to the value vector of the dimension *L × M ×* 64 (M: the number of sequences in the MSA). In order to extract more important amino acids in each column of MSA, we also apply the column attention to the input MSA embedding vector by using the linear transformation and softmax function. The attention weight has the dimension of *L × M ×* 8. The final output of the input embedding component is the multiplication of the column attention weights and the input MSA embedding vector with the dimension of *L* × 512. In order to maintain the order of the sequence, we add the positional encoding to the final embeddings. The input embedding vector is further processed by the self-attention component composed of a stack of *N* = 6 identical sub-components to generate the output of the encoder. Each sub-component is composed of a multi-head self-attention layer(*numberofheads* = 8) and a fully connected feed-forward layer. For each layer, there is a residual connection[39] around it, followed by the layer normalization[40] and one dropout operation[41] with dropout rate = 0.1.

Similar to the encoder, the decoder also consists of the embedding component and two attention components. It starts from the output tokens (i.e., true 3-state secondary structure in the training phase or predicted secondary structure in the inference phase) of the previous positions in a target protein, which are converted to an embedding vector of the dimension *L ×* 512. A sinusoidal position encoding vector is added on top of the embedding vector, which is then fed into the first self-attention component, followed by the cross-attention component. The first self-attention component consists of a stack of *N* = 6 identical layers, similar to the attention component in the encoder except that it applies the masked self-attention mechanism to ensure that the prediction for the current position only depends on the output tokens prior to this position in the training phase. In the inference phase, the self-attention component in the decoder also considers only the previously generated secondary structure states when generating the next. Both the output of the self-attention component in the decoder and the output of the encoder stack are fed into the crossattention decoder component composed of N = 6 identical sub-components, each consisting of a multi-head self-attention layer(*numofheads* = 8) and a fully connected feed-forward layer. The output of the decoder stack is converted to the 3-state protein secondary structure prediction through the linear transformation and softmax function.

We use Adam[42] as the optimizer with the following parameters: *β*1 = 0.9, *β*2 = 0.999, warmup steps = 4000, and weight decay = 0.001. We set the batch size to 30 and use the Kullback-Leibler divergence loss as the loss function to train TransPross. We set the number of training epochs to 100 with the early stopping to reduce overfitting. If there is no improvement in the validation loss for five consecutive epochs, the training stops. We train through 10-fold cross validation. Five TransPross best models have been selected and the final output is the ensemble of the five models.

### 3.3 Evaluation metrics

We compare TransPross with a state-of-the-art profile-based protein secondary structure predictor - PSIPRED and a single sequence predictor - SPOT-1D. We evaluate the prediction performance in terms of the Q3 accuracy score, commonly used in the secondary structure prediction. The Q3 accuracy score is defined as the percentage of the correctly predicted secondary structures in three categories (strand E, helix H, and coil C)[43].

## 4 Conclusion

In this work, we introduce a novel transformer method based on the language model (TransPross) for protein secondary structure prediction. It takes a MSA of a protein as input and automatically encodes it as internal features for a decoder to predict 3-class secondary structures of the protein. By using the column attention, TransPross is able to assign attention weights in each column corresponding to each residue position in the protein and extracts the evolutionary patterns to predict the per-residue secondary structure based on all the inter-dependencies over columns and rows across the whole MSA. In addition to achieving the state-of-the-art performance, TransPross can take a user-provided MSA from any source to generate secondary structure prediction quickly, making it a convenient tool for users to predict secondary structures at a large scale. TransPross can also be used by developers to create more deep learning language and transformer models to improve protein secondary structure prediction.

## Declarations

### Consent for publication

All the authors approve the manuscript for publication.

### Availability of data and materials

The data is available at https://zenodo.org/record/6762376#.Ys4St5PMKsA.

### Code availability

The source code is available at https://github.com/BioinfoMachineLearning/TransPro.

### Authors’ contributions

JC conceived the project. TW and JC designed the method. TW implemented the method, performed the experiment, and collected the data. TW and JC analyzed the data. WC reviewed and summarized some related methods. TW and JC wrote the manuscript.

